## Supplemental Figure 1 for "Downregulation of *Let-7* miRNA promotes Tc17 differentiation and emphysema via de-repression of RORγt"

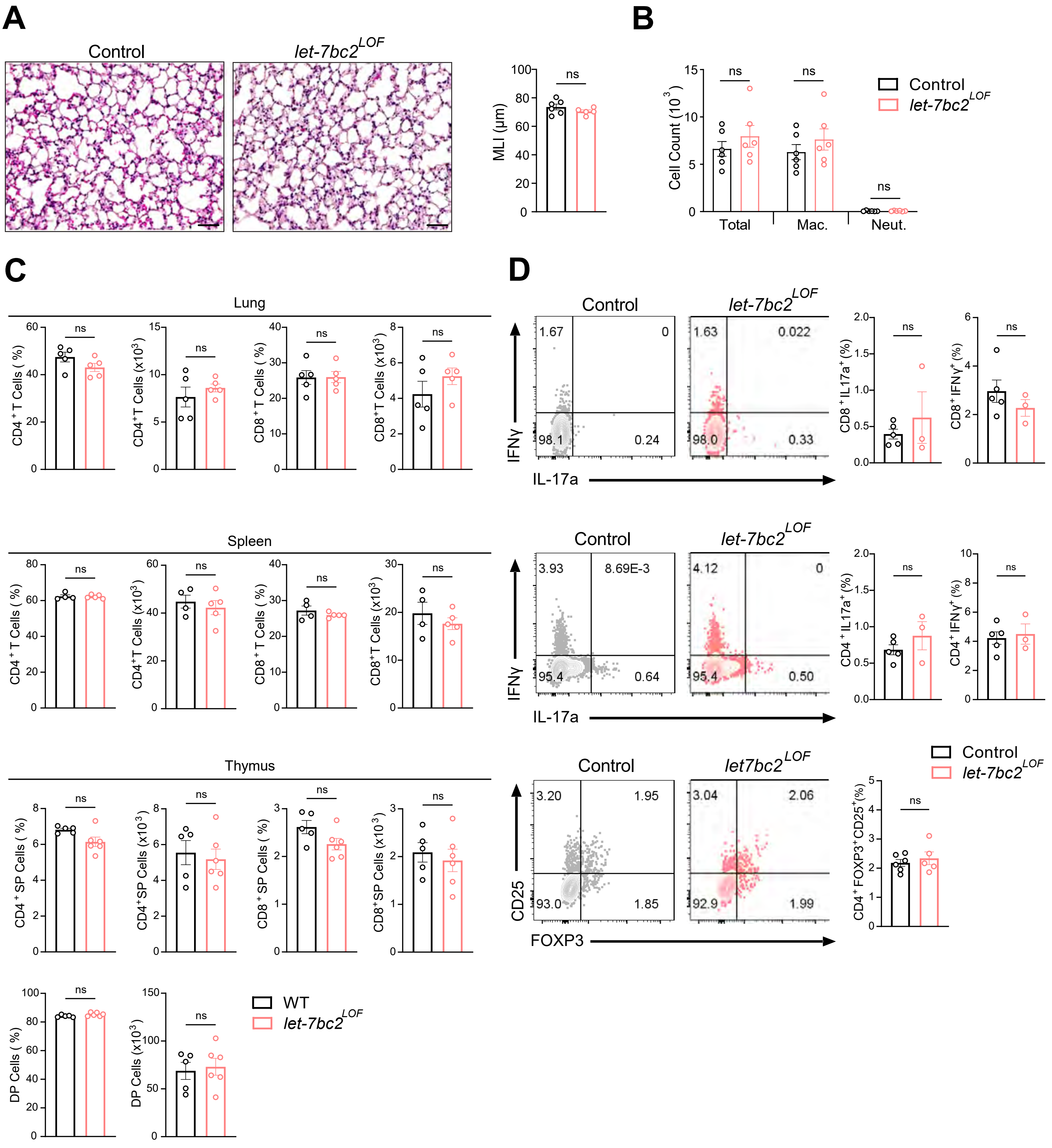


**Figure 2-figure supplement 1. T cell-specific deletion of the *let-7bc2*-cluster does not promote lung inflammation or pathology with moderate aging.** (A) Representative H&E-stained lung sections from Control and *let-7bc2*$^{LOF}$ naive mice aged to one year (x20 magnification; scale bars, 50μm) with MLI measurement of lung morphometry (n=5-6 per group). (B) Total and differential cell count from bronchoalveolar lavage (BAL) fluid from control and *let-7bc2*$^{LOF}$ naive mice (n=6 per group; Mac. = macrophages, Neut. = neutrophils). (C) Flow cytometric analysis of CD4$^+$, CD8$^+$, or DP T cells from the lungs, spleen, and thymus of control and *let-7bc2*$^{LOF}$ mice at steady-state (n=4-6 per group). (D) Immunophenotyping of Tc17, Tc1, Th17, Th1, and Tregs from lungs of naïve control and *let-7bc2*$^{LOF}$ mice (n=3-5 per group). Data are representative of three independent experiments and displayed as mean±SEM using student's t-test.
