## Supplemental Figure 2 for "Downregulation of *Let-7* miRNA promotes Tc17 differentiation and emphysema via de-repression of RORγt"

**A**

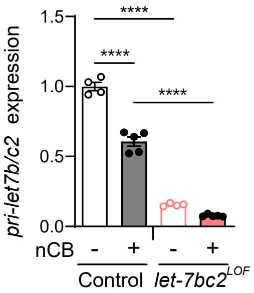


**Figure 2-figure supplement 2. *Let-7bc2* cluster expression in lung CD8$^+$ T cells of naive and nCB-exposed mice.** (A) QPCR analysis of *pri-let7b/c2* from sorted lung CD8$^+$ T cells of PBS vehicle or nCB treated control and *let-7bc2$^{LOF}$* mice (n=3-6 per group). Data are representative of two independent experiments and displayed as mean±SEM using two-way ANOVA with *post-hoc* Tukey correction. ****p<0.0001.
