## Supplemental Figure 3 for "Downregulation of *Let-7* miRNA promotes Tc17 differentiation and emphysema via de-repression of RORγt"

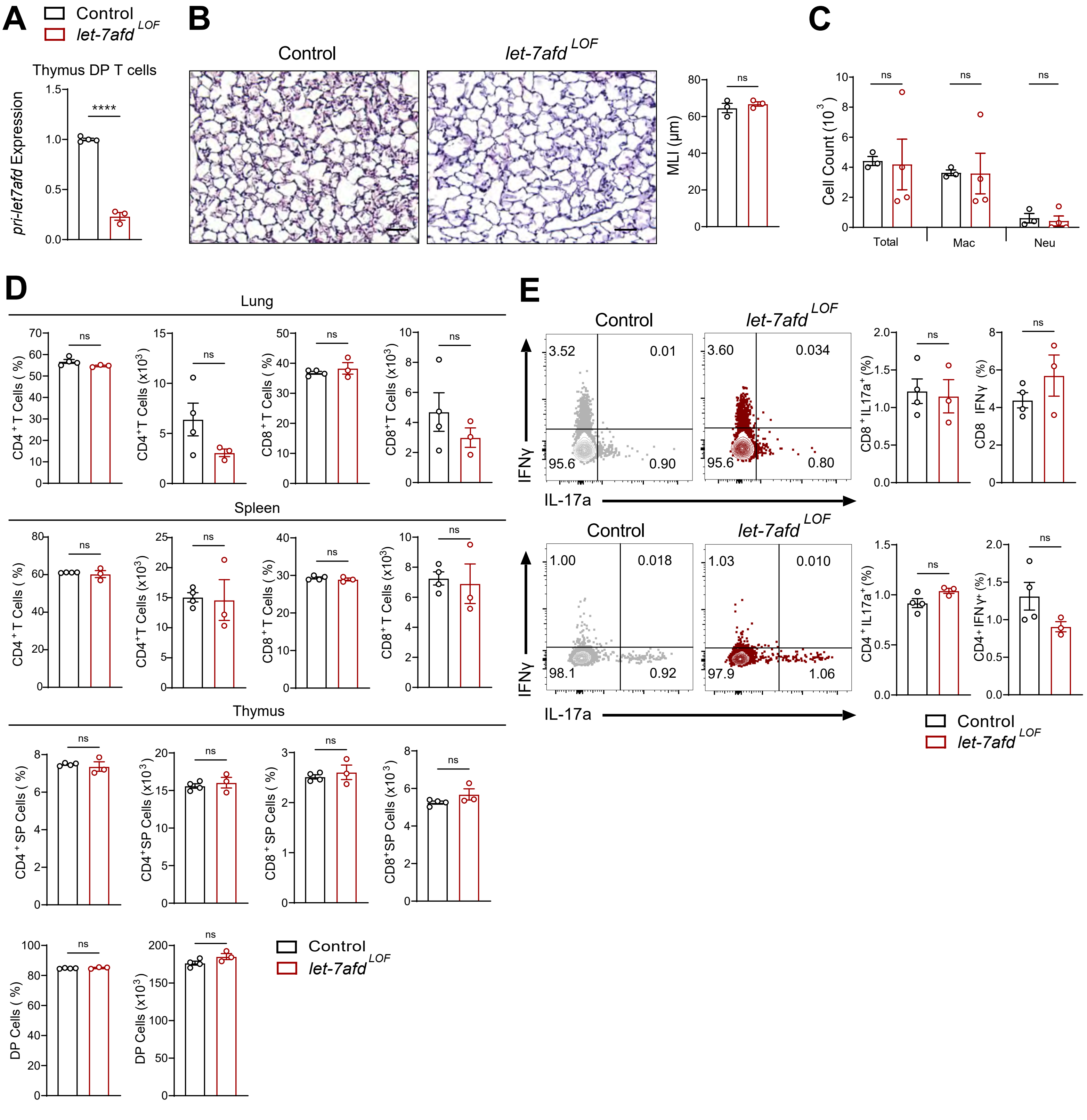

**Figure 4-figure supplement 1. T cell-specific deletion of the *let-7afd*-cluster does not promote lung inflammation or pathology with moderate aging.** (A) QPCR analysis of *pri-let-7a1/f1/d* from flow-sorted live, TCRβ⁺, CD4⁺CD8 double-positive (DP) thymocytes of control and *let-7afd*$^{LOF}$ mice (n=3-4 per group). (B) Representative H&E-stained lung sections from control and *let-7afd*$^{LOF}$ naive mice aged to 6 months (x20 magnification; scale bars, 50µm) with MLI measurement of lung morphometry (n=3 per group). (C) Total and differential cell count from bronchoalveolar lavage (BAL) fluid from control and *let-7afd*$^{LOF}$ naive mice (n=3-4 per group; Mac. = macrophages, Neut. = neutrophils). (D) Flow cytometric analysis of CD4⁺, CD8⁺, or DP T cells from the lungs, spleen, and thymus of control and *let-7afd*$^{LOF}$ mice at steady-state (n=3-4 per group). (E) Immunophenotyping of Tc17, Tc1, Th17, and Th1 cell from lungs of naïve control and *let-7afd*$^{LOF}$ mice (n=3-4 per group). Data are representative of two independent experiments and displayed as mean±SEM using student'st-test. ****p<0.0001.
