## Supplemental Figure 4 for "Downregulation of *Let-7* miRNA promotes Tc17 differentiation and emphysema via de-repression of RORγt"

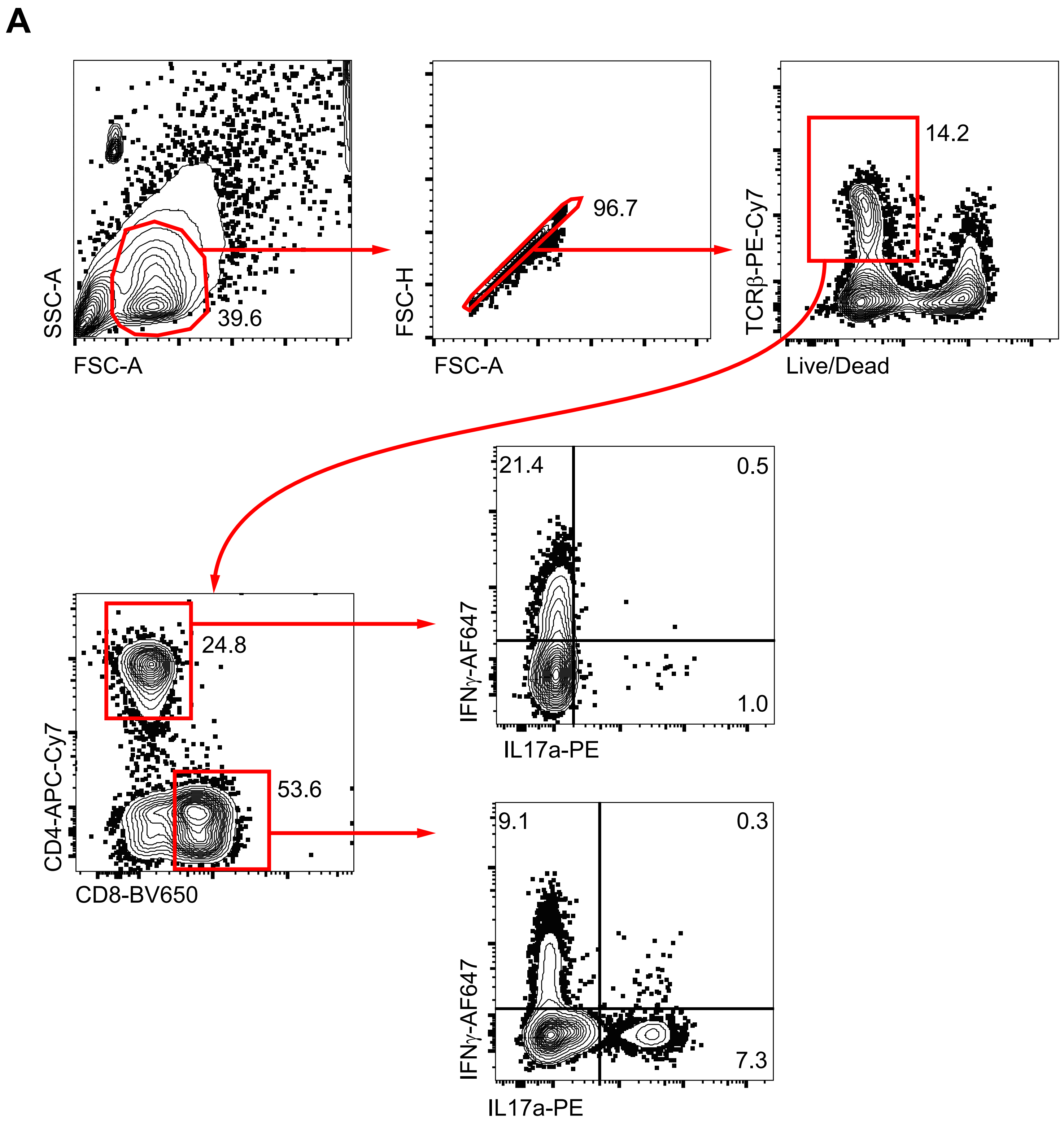


**Supplementary Figure 3.** (A) Representative flow gating strategy for the identification of Th1/Th17 and Tc1/Tc17 T cell subsets in mouse lungs of wild-type control mouse.
