## Supplemental Table 1 for "Downregulation of *Let-7* miRNA promotes Tc17 differentiation and emphysema via de-repression of RORγt"

**Supplementary Table 1: Genotyping Primers and Duplexes for Luciferase Assay**

**Genotyping Primers**

| **Name** | **Sequence** | **Description** |
|---|---|---|
| BC-lox2-F | 5'-GGACATGAGATCGCCAACCA-3' | *let-7bc2*-cluster floxed allele<br>Forward primer for genotyping |
| BC-lox2-R | 5'-TGGAAGCCAGTACTGTGCTC-3' | *let-7bc2*-cluster floxed allele<br>Reverse primer for genotyping |
| AFD-lox2-F | 5'-GTTTTCTGAGGTGTGGGAGGTA-3' | *let-7afd*-cluster floxed allele<br>Forward primer for genotyping |
| AFD-lox2-R | 5'-AGTGGGATAGAAGGATCTCAGG-3' | *let-7afd*-cluster floxed allele<br>Reverse primer for genotyping |

**Dharmacon Duplexes**

| **Name** | **Sequence** |
|---|---|
| *hsa-let-7b-5p* | 5'-UGAGGUAGUAGGUUGUGUGGUU-3' |
| Control (*cel-miR-67-3p*) | 5'-UCACAACCUCCUAGAAAGAGUAGA-3' |
