## Supplemental Table 2 for "Downregulation of *Let-7* miRNA promotes Tc17 differentiation and emphysema via de-repression of RORγt"

**Supplementary Table 2. Demographics of subjects by emphysema severity**

| Emphysema Severity Score | = 0 | = 1 | = 2 | = 3 |
|---|---|---|---|---|
| Sex, n (%) | | | | |
| Male | 2 (100) | 5 (100) | 5 (100) | 6 (85.7) |
| Female | 0 (0) | 0 (0) | 0 (0) | 1 (14.3) |
| Age | 74.0 ± 4.2 | 66.0 ± 7.6 | 65.2 ± 4.6 | 67.1 ± 3.9 |
| Race, n (%) | | | | |
| Caucasian | 2 (100) | 5 (100) | 4 (80) | 5 (71.4) |
| African American | 0 (0) | 0 (0) | 1 (20) | 2 (28.6) |
| Current Smoker, n (%) | | | | |
| Yes | 0 (0) | 2 (40) | 3 (60) | 4 (57.1) |
| No | 2 (100) | 3 (60) | 2 (40) | 3 (42.9) |
| $FEV_1$ % | 57.0 ± 17.0 | 68.8 ± 12.3 | 74.2 ± 13.9 | 80.7 ± 9.5 |
| $FEV_1$/FVC % | 68.5 ± 12.0 | 66.6 ± 5.7 | 70.3 ± 9.4 | 60.3 ± 6.7 |

Mean ± standard deviation is shown unless otherwise stated.
